# A Coiled-Coil-Based Design Strategy for the Thermostabilization of G-Protein-Coupled Receptors

**DOI:** 10.1101/2022.03.28.485961

**Authors:** Marwa Amer, Daniel Frey, Xiaodan Li, Richard A. Kammerer

**Author notes:** Corresponding author: Richard A. Kammerer, Laboratory of Biomolecular Research, Division of Biology and Chemistry, Paul Scherrer Institute, 5232 Villigen PSI, Switzerland.

## Abstract

The most common methods for generating crystallizable GPCRs are scanning alanine mutagenesis and fusion to crystallization-facilitating partner proteins. The major goal of our work was to create a new GPCR tool that would provide receptor stability and additional soluble surface for crystallization. Towards this aim, we selected the two-stranded antiparallel coiled coil as a domain fold that satisfies both criteria. A selection of antiparallel coiled coils was used for structure-guided substitution of intracellular loop of the β3 adrenergic receptor. Unexpectedly, only the two GPCR variants containing thermostable coiled coils were expressed. We showed that one GPCR chimera is stable upon purification in detergent, retains ligand-binding properties, and can be crystallized. However, the quality of the crystals was not suitable for structure determination. To supply additional surface for promoting crystal contacts, we replaced in a structure-based approach the loop of the antiparallel coiled coil by T4L. Although expression is currently not suitable for structural work, we found that the engineered GPCR is even more stable than the coiled-coil variant. Our approach should be of interest for applications that benefit from stable GPCRs.

## INTRODUCTION

G-protein-coupled receptors (GPCRs) are characterized by a seven-transmembrane helix topology and represent the largest membrane protein family [1]. GPCRs play fundamental roles in almost all physiological and pathological processes by responding to a variety of extracellular signals, including photons, small molecules, peptides, and proteins. These signals cause conformational changes in the GPCR and lead to the activation of associated G proteins that regulate central downstream signaling pathways [2, 3]. GPCRs are the target of approximately 30% of all approved drugs on the market and are therefore of enormous medical and commercial interest [4-6]. Despite AlphaFold2 and RoseTTAfold [7, 8] that both can predict apo-structures to a high degree of accuracy, there is a growing demand on high-resolution structures of GPCR/ligand complexes for structure-based drug design. However, generating GPCRs suitable for X-ray structural studies is a challenging subject because of their poor expression, conformational heterogeneity, and stability that affects purification and crystallization [9]. Accordingly, the determination of available GPCR crystal structures was the result of extensive modifications that often required a combination of several of the following protein engineering approaches: 1) truncation of unstructured N- and/or C-termini and/or loops, 2) scanning alanine mutagenesis (SAM) 3) application of fusion proteins, such as bacteriophage T4 lysozyme (T4L), thermostabilized cytochrome b562RIL (BRIL) and others, 4) removal of post-translation modification sites and 5) stabilization by antibodies and nanobodies [9].

The most common methods used for crystallizing GPCRs are SAM [10] and the fusion protein approach [11]. The rationale behind SAM is to create stable and conformation-specific GPCRs while maintaining their pharmacological activity. Systematically, single amino acid mutants of a GPCR of interest are generated by substituting each amino acid in the sequence with Ala (Gly if the wild-type residue is an Ala). Typically, the most stabilizing mutations are then combined until a mutant with the desired stability is obtained. This method was for example successfully employed for solving the structure of the turkey β1 adrenergic receptor [12] and the C-C chemokine receptor type 9 [13]. A major disadvantage of SAM is that the method is rather labor-intensive. Moreover, the stabilized GPCR might still require further protein engineering to generate the necessary soluble surface area needed for crystallization in detergent.

Extension of the relatively small polar surface area of GPCRs that is available for forming crystal lattice contacts is the rationale behind the fusion protein engineering approach [11]. Typically, partner proteins such as T4L, bRIL and others are fused to the truncated N-terminus of a GPCR or replace the unstructured intracellular loop 3 (ICL3) [9]. Using this approach, the crystal structures of more than 40 receptors were determined, including those of the β2 adrenergic receptor, the histamine H1 receptor [14], the dopamine D3 receptor [15], the chemokine CXCR4 receptor [16] and the human CC chemokine receptor 7 [17]. Not all GPCRs are amenable to fusion protein engineering, and it appears that the approach is suitable for GPCRs only that when bound to a stabilizing ligand are stable upon detergent solubilization from their membrane environment.

Based on the available crystal structures, it seems that a combination of both SAM and the fusion protein approach would be the most promising strategy for the efficient crystallization of GPCRs. The major goal of our work was to create a new GPCR tool that would provide receptor stability and additional crystallizable surface at the same time. Toward our aim, we selected the α-helical coiled coil as an ideal candidate because coiled coils are generally very soluble and can fold into very stable structures. Furthermore, short coiled coils typically can be easily crystallized and resulting crystal structures are frequently determined at high resolution. The left-handed coiled coil is probably the most widespread subunit oligomerization motif found in proteins [18-21]. It consists of two or more amphipathic α-helices that “coil” around each other in a left-handed supertwist. It characterized by a heptad-repeat sequence of seven amino-acid residues denoted *[abcdefg]*_*n*_ (Fig.1 *B*) with a 3,4-hydrophobic repeat of mostly non-polar amino acids at positions *a* and *d* [22, 23]. Interactions between the *a* and *d* core residues and its flanking *e* and *g* positions determine the stability of a coiled coil, the number of strands it consists of, the parallel or antiparallel orientation of α-helices, and the homo- or heterotypic association of subunits.

A selection of different two-stranded antiparallel coiled-coil structures, in which the same polypeptide chain folds back on itself, were used to replace ICL3 of the β3 adrenergic receptor (β3AR). Expression, solubilization in detergent, analytical and fluorescence size exclusion chromatography, and binding studies were performed, to assess the effect of the coiled coils on the functionality and stability of the chimeric receptor.

## RESULTS AND DISCUSSION

### Design rationale of engineered GPCR variants

The design rationale of our GPCR variants is based on the observation that at the cytoplasmic site in the crystal structure of β2AR, transmembrane helices 5 and 6 are interacting over a short stretch of residues in a way that is very similar to an antiparallel coiled coil [24] (PDB code: 2RH1). Specifically, residues Ala226 and Phe223 of helix 5 that can be assigned the hydrophobic heptad-repeat positions *d* and *a*, respectively, interact with His269 and Leu272 at positions *a’* and *d’*, respectively, of helix 6 (Fig. 1A). β3AR was selected because it is the closest relative of β2AR, for which no high-resolution structure is available. The idea behind the design was to replace ICL3 of β3AR by a series of antiparallel coiled coils and thereby extend the sequence corresponding to coiled-coil-like structure seen in β2AR (Fig. 1B and C). The coiled coil would therefore act like a clamp on β3AR transmembrane helices 5 and 6 that would hopefully stabilize the entire GPCR and provide additional soluble surface for crystallization. The coiled coils were inserted into ICL3 between Ala231 and Glu286 (Fig. 1B).

**Figure 1.**
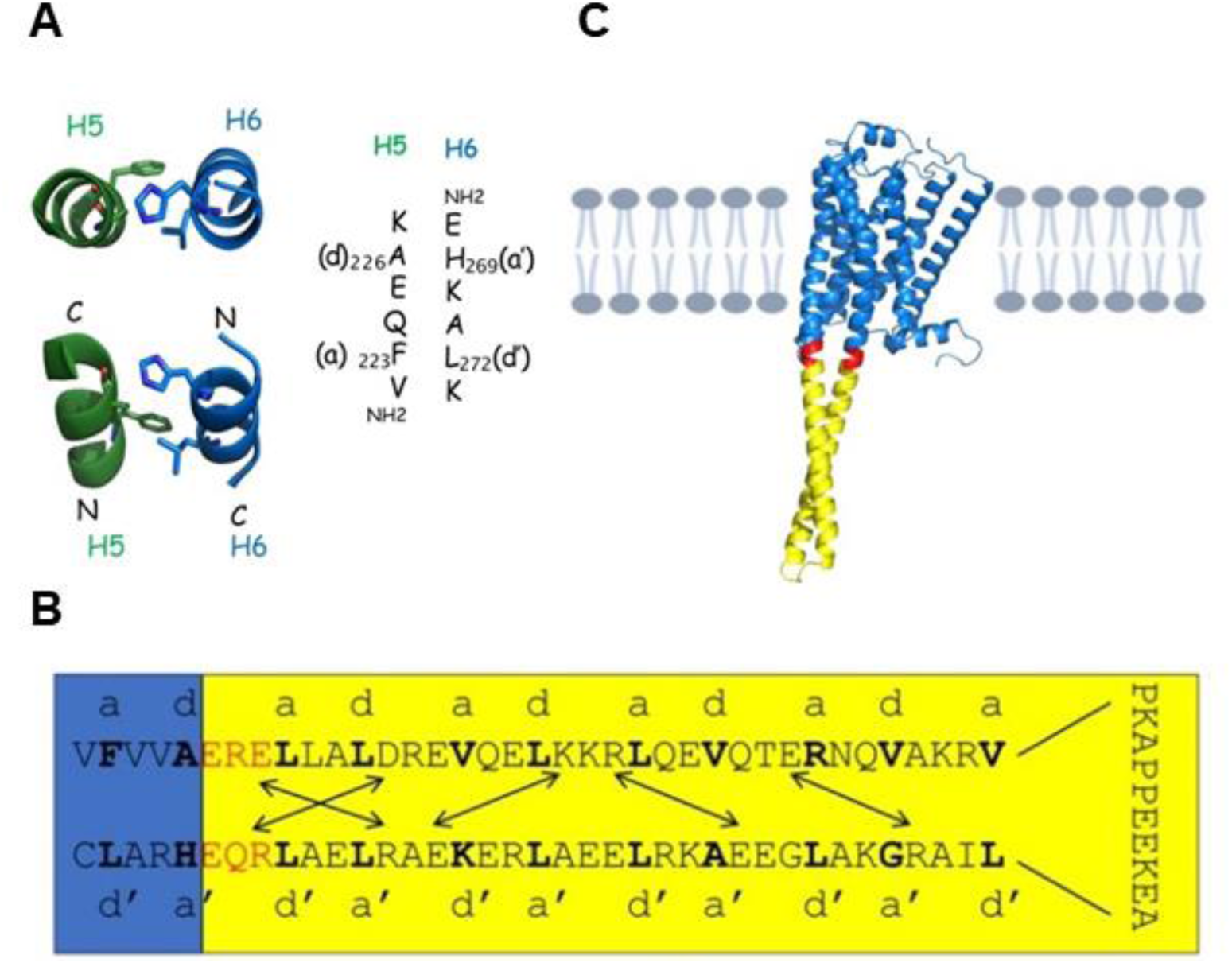
Design rational of β3AR-coiled-coil chimeras. (**A**) Top (upper panel) and side view (lower panel) of ribbon representation of crystal structure of β2-adrenergic receptor transmembrane helices 5 (green) and 6 (blue) that interact in n antiparallel a coiled-coil like manner at the cytosolic side (PDB code: 2RH1) [24]. Interacting amino-acid residues are shown as sticks. Sequences of the interacting segments of transmembrane helices 5 and 6 are shown in the right panel. Residues at the hydrophobic heptad-repeat positions a, a’, d and d’ are numbered according to their position in the wild-type protein. N- and C-termini are indicated. (**B**) Example of chimeric β3AR. In-register heptad-repeat fusion of the antiparallel coiled coil of the *Thermus thermophilus* seryl-tRNA synthetase (SRS, cc3.1) (yellow) and the human β3-adrenergic receptor (blue). Introduction of three amino-acid residues at the junction (red) was necessary to obtain a continuous heptad-repeat pattern. Amino-acid residues at the e’ and f positions of the junction were chosen to introduce two additional attractive salt bridges into cc3.1, to potentially further stabilize the protein. The heptad repeat pattern is indicated. Potential salt bridges are indicated by arrows. (**C**) β3AR-cc3.1 model. β3AR is shown in blue, the cc3.1 coiled coil in yellow and residues at the junction in red. The model was generated by AlphaFold2 [7].

Coiled-coil positions *e, e’, g* and *g’* that flank the hydrophobic *a, a’, d* and *d’* positions are frequently occupied by charged residues that form *g* to *g’* and *e* to *e’* type salt bridges that can further stabilize the structure [25, 26]. To this end, amino-acid residues at the junction were chosen to optimize attractive electrostatic interactions between helices (Fig. 1B).

Extensive screening of the RCSB Protein Data Bank and the literature was performed, to identify suitable antiparallel coiled-coil candidates for substituting ICL3 of β3AR. Based on their characteristics, six antiparallel coiled coils were selected (Fig. S1 and table S1). cc1 is derived from a bacteriophage serine integrase that plays a key role in the integration of the viral genome through self-interactions between the coiled-coil domains [27]. Based on its small size (three heptad repeats), the coiled coil is not expected to affect G-protein binding of the β3AR chimera. cc2 originates from a viral nucleocapsid protein [28] and like cc1 can interact with itself. The capability to self-interact makes cc1 and cc2 promising candidates to establish potential crystal contacts between chimeric β3AR-cc molecules. Coiled coils from thermostable proteins are potentially extremely stable and attractive candidates to stabilize β3AR. Towards this aim, the antiparallel coiled-coil domains of seryl-tRNA synthetase (SRS) from *Thermus thermophilus* (cc3.1) and *Pyrolobus fumarrii* (cc3.2), bacteria that can grow in extreme temperature conditions of up to 85°C and 122°C, respectively, were selected [29]. Although high-resolution structures of the isolated thermostable coiled-coils are not available for the proteins, the domain from *E. coli* has been characterized in detail [30]. Coiled-coil candidates with defined additional α-helical structures instead of the loop connecting the two helices were also selected because they provide a larger surface for crystallization than classical antiparallel coiled coils. cc4 is derived from the pore-forming toxin YaxAB [31] and cc5 represents the microtubule-binding domain from dynein [32].

Covalent connection of helices by a disulfide bond at its N- or C-terminus is a common approach to stabilize a coiled-coil structure. Because some of the isolated coiled-coil candidates are potentially not very stable, a second series of β3AR chimeras harboring an intramolecular disulfide bond, termed βAR3-cc_SS, was designed to increase their stability [33] (Fig. S1 and table S1).

Based on the secondary structure prediction, truncation of the predicted unstructured N- and C-terminus of human β3AR was carried out. Specifically, the 25 N-terminal residues (Pro3 to Thr27) that contain two potential N-glycosylation sites (amino-acid residues Asn8 and Asn26) and the C-terminal 40 residues after the palmitoylation site (Pro369 to Ser408) were deleted.

Residue substitutions were identified that thermostabilized the turkey β1AR. The combination mutant was significantly more stable than the native protein when solubilized in dodecylmaltoside (DDM) and in short chain detergents, which allowed its crystallization and structure determination. Because it was shown that the mutations could be transferred from β1AR to β2AR [34], the β3AR sequence was further modified by introducing these mutations: Glu36Ala, Met86Val, Ile125Val, Glu126Trp, Tyr234Ala, Phe341Met, and Tyr346Leu [35].

### Only β3AR-cc chimeras fused to thermostable coiled coils exhibit expression

Three different cell lines, HEK293S GnTI^-^, T-REx-293, T-REx-CHO, were screened for the expression of the 12 β3AR chimeras. Only β3AR-cc3.1 and β3AR-cc3.2 showed reasonable expression levels in T-REx-293 cells. The expression level of β3AR-cc3.1 was higher compared to the one of β3AR-cc3.2. Instead, the other tested variants did not express in any of the cell lines. These results suggest that the stability of GPCRs is an important factor for expression, a hypothesis that is consistent with that we were unable to express wild-type β3AR as a control. They furthermore demonstrate that thermostable coiled coils are promising fusion candidates for GPCR stabilization and that the stabilizing mutations identified for β1AR had no stabilizing effect on β3AR.

The observation that the disulfide-linked variants of the β3AR-cc3.1 and β3AR-cc3.2 chimeras did not express can be rationally explained by destabilization of the coiled coils upon introducing Cys residues. Hydrophobic amino acids like Leu, Ile or Val at heptad repeat *a* and *d* positions are mainly responsible for the stability of a coiled coil and substitution of such a residue usually leads to a significant destabilization of the structure [36, 37].

As a result of its higher expression, we focused on β3AR-cc3.1 and generated stable cell lines expressing chimeric protein. Notably, addition of both tetracycline (2 μg/ml) and sodium butyrate (5mM) resulted in a three-fold increase in the expression efficiency.

### β3AR-cc3.1 is stable upon solubilization and purification in detergent

To identify the best detergent for solubilization, a screen of representative detergents from different families covering a wide range of chemical properties was carried out. To this aim, (n-dodecyl-β-d-maltopyranoside (DDM) (maltoside detergent group), lauryl maltose neopentyl glycol (LMNG) (NG class detergent group), undecanoyl-n-hydroxyethylglucamide (HEGA11) (HEGA detergent group), n-decyl-β-d-thiomaltoside (DDTM) (thio maltoside detergent group), n-dodecylphosphocholine (FC12) (lipid-like detergent group), 5-cyclohexyl-1-pentyl-β-d-maltoside (CYMAL7) (CYMAL detergent group) and n-dodecyl-β-d-glucopyranoside (DDG) (glucoside detergent group) and n-decyl-β-d-glucopyranoside (DTG) (thio glucoside detergent group) were tested. Each detergent was assessed for its ability to solubilize β3AR-cc3.1 at a final concentration of 1% (w/v), a concentration that is at least 100 times above the CMC values of the tested detergents. Western blot analysis using an anti-FLAG monoclonal antibody demonstrated that all the detergents efficiently solubilized β3AR-cc3.1 (Fig. 2A).

**Figure 2:**
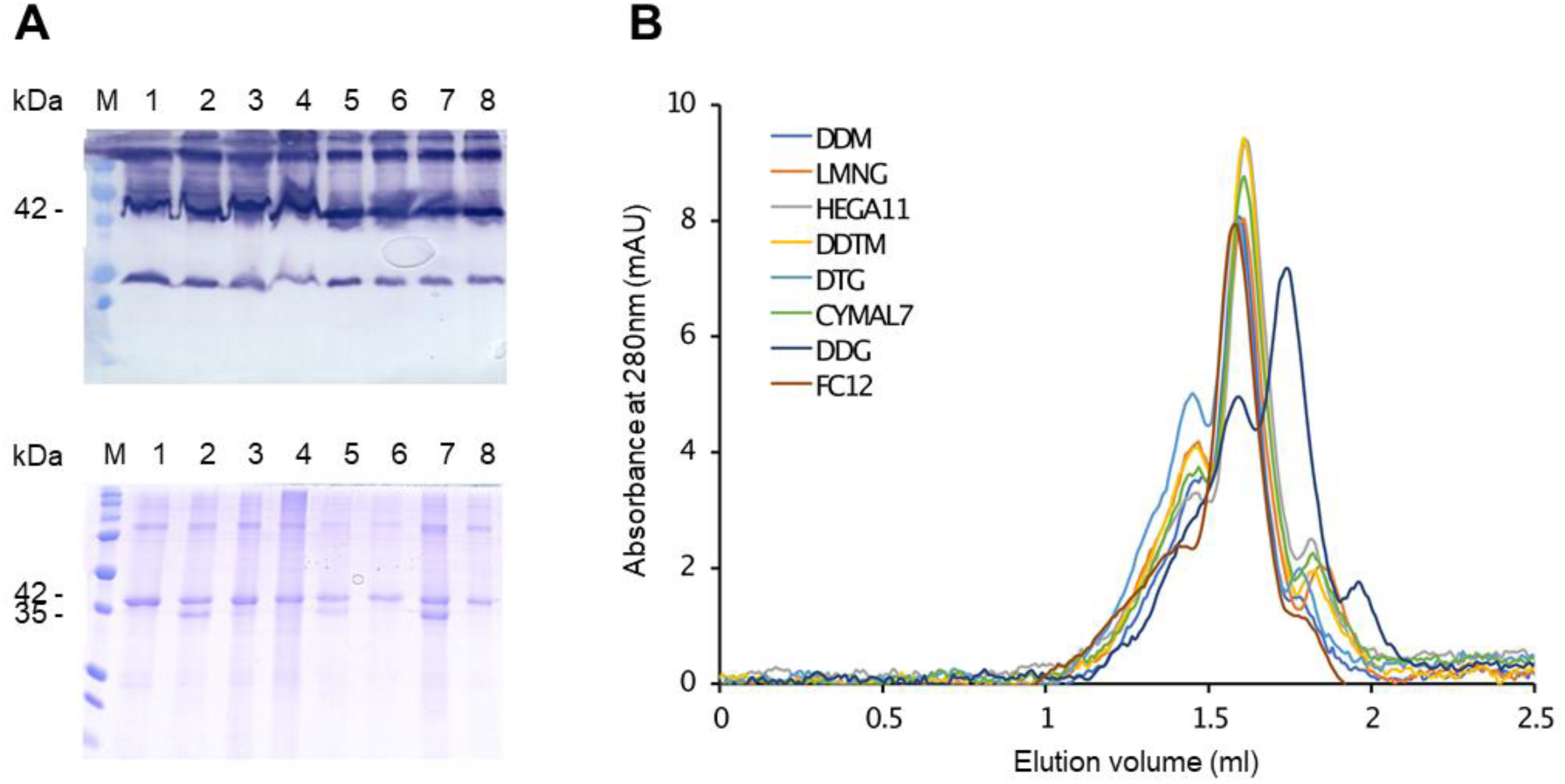
Solubilization, purification and monodispersity of β3AR-cc3.1. (**A**) Western blot (upper panel) and SDS-PAGE (lower panel) analysis of solubilization and purification of β3AR-cc3.1. All used detergents efficiently solubilized β3AR-cc3.1. The faster migrating band observed for some detergents could represent a SDS-resistant conformation of β3AR-cc3.1. Lane 1, DDM; lane 2, LMNG; lane 3, HEGA11; lane 4, DDTM; lane 5, DTG; lane 6, CYMAL7; lane 7, DDG, lane 8, FC12. The migration of marker proteins (M) is shown. (**B**) Analytical size exclusion chromatography for β3AR-cc3.1 purified in different detergents. The protein is eluting at a volume that corresponds to a monomer. Concerning monodispersity, the best detergent for the purification was DDM/CHS followed by FC12 and HEGA11.

Next, a small-scale purification of β3AR-cc3.1 using StrepTrap Sepharose beads was performed for each detergent as described in the Materials and Methods section. SDS-PAGE analysis demonstrated that β3AR-cc3.1 could be efficiently purified with all the tested detergents. For most detergents, a single band of β3AR-cc3.1 migrating at approximately 42kDa was detected, but for LMNG, DTG and DDG an additional band of approximately 35kDa was observed (Fig. 2A). The faster migrating band could represent a SDS-resistant conformation of β3AR-cc3.1 because many membrane proteins migrate faster on SDS-PAGE than their predicted molecular mass [38].

Therefore, analytical size exclusion chromatography was used in a next step to assess the monodispersity of the purified protein samples. As can be seen in figure 2B, a combination of DDM and cholesterol hemisuccinate (CHS) followed by FC12 and HEGA11 were the best detergents in terms of β3AR-cc3.1 monodispersity. Our findings are consistent with the observation that DDM/CHS was previously very successfully used for the purification of several GPCRs [39]. In the following we focused on the three best detergents identified in our experiments.

### The thermostable SRS coiled coil significantly stabilizes β3AR

Next, we assessed the thermal stability of β3AR-cc3.1 using the thiol-specific probe, 7-diethylamino-3-(4-maleimidophenyl)-4-methylcoumarin (CPM) [40]. The assay measures the fluorescence emission of CPM upon forming a covalent bond with the side chain of a free Cys. The free cysteine becomes more readily accessible upon protein thermal denaturation. β3AR-cc3.1 contains 14 cysteine residues. The CPM measurements were carried out in the range of 25°C to 90°C using a ramping rate of 2°C/min to warrant equilibrium during unfolding without compromising the integrity of CPM. The thermal stability of β3AR-cc3.1 was tested with different ligands (antagonists: SR59230A, L748337 and carvedilol; agonist: carazolol), in Tris-HCl and HEPES buffers, and at different protein concentrations. For β3AR-cc3.1, a melting temperature (T_m_) value of 64°C was obtained in DDM/CHS and carvedilol (Fig. 3). In comparison, the T_m_s of thermostabilized GPCRs were in the range of 45–55°C [41, 42], demonstrating that our approach is well suited to stabilize GPCRs. As expected, the stability of β3AR-cc3.1 was dependent on the type of detergent that was used. The T_m_ of the purified β3AR-cc3.1 decreased by 6°C and 10°C in the presence of HEGA11 and FC12, respectively (Fig. 3).

**Figure 3:**
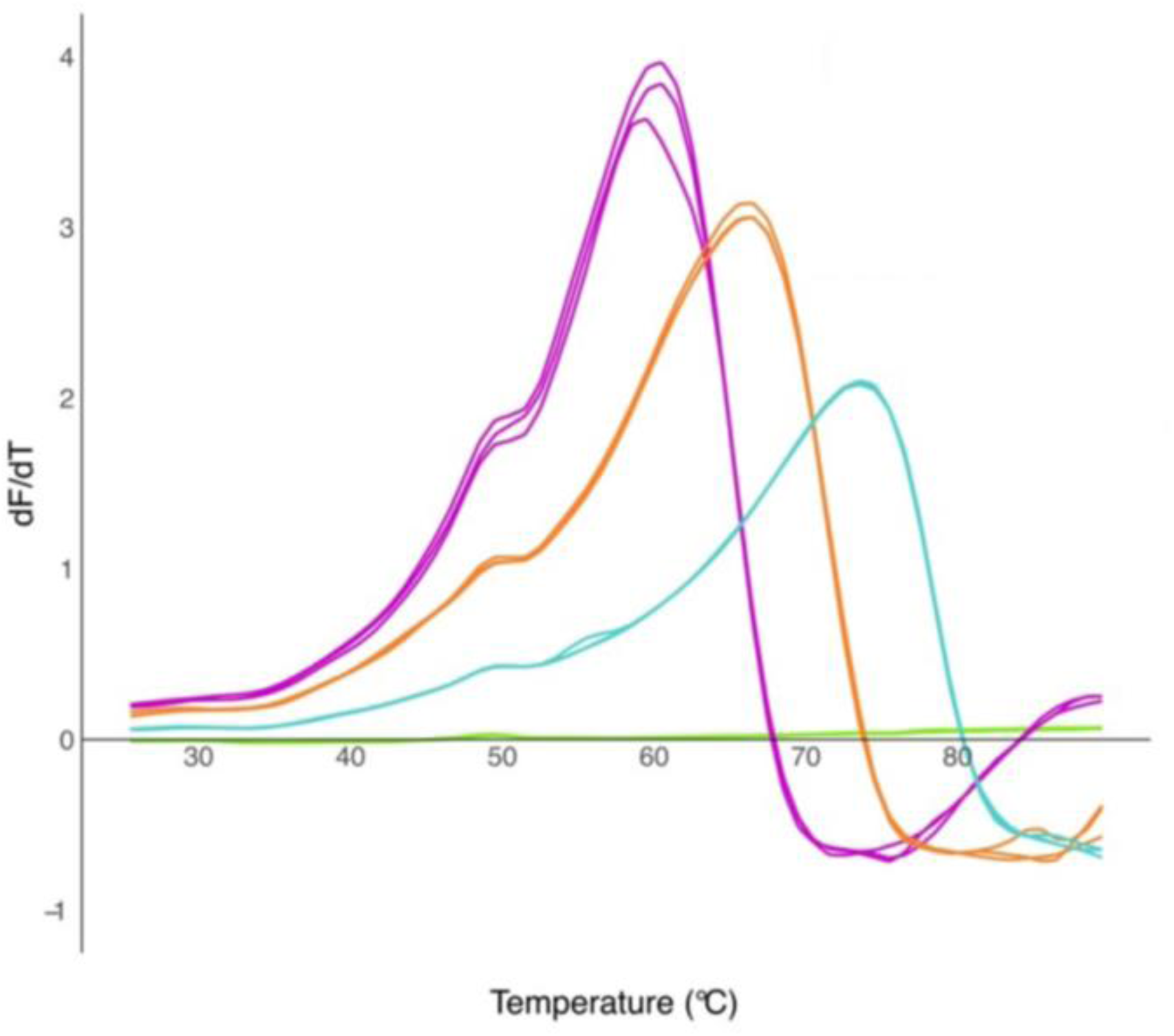
Thermal stability of β3AR-cc3.1. Replicate CPM measurements of β3AR-cc3.1 (15µg) bound to carvedilol in different detergents. The apparent melting temperature of β3AR-cc3.1 in FC12 (purple) was 55 °C, 57 °C in HEGA11 (orange) and 65 °C in DDM/CHS (cyan). The light green curves represent measurements of the buffer without any protein.

### β3AR-cc3.1 maintains ligand-binding activity

TM helices 5 and 6 play important roles for ligand binding and receptor activation [3]. Because our design approach to link TM helices 5 and 6 by a thermostable antiparallel coiled coil probably limits their conformational flexibility, it was important to assess the ligand binding properties of β3AR-cc3.1. To this end, saturation-binding assays were carried out with isolated HEK-239S-GnTI^-^membranes expressing β3AR-cc3.1. For these experiments, the antagonist [3H] dihydroalprenolol (DHA) was used for which a K_D_ of ∼100 nM for the binding to β3AR has been reported [43]. Although DHA binding to β3AR-cc3.1 was not fully saturated at 300nM, a K_D_ value of 150 nM was estimated (Fig. 4). This value is similar to the previously reported K_D_, demonstrating that the engineered GPCR is still capable of binding to the antagonist ligand.

**Figure 4.**
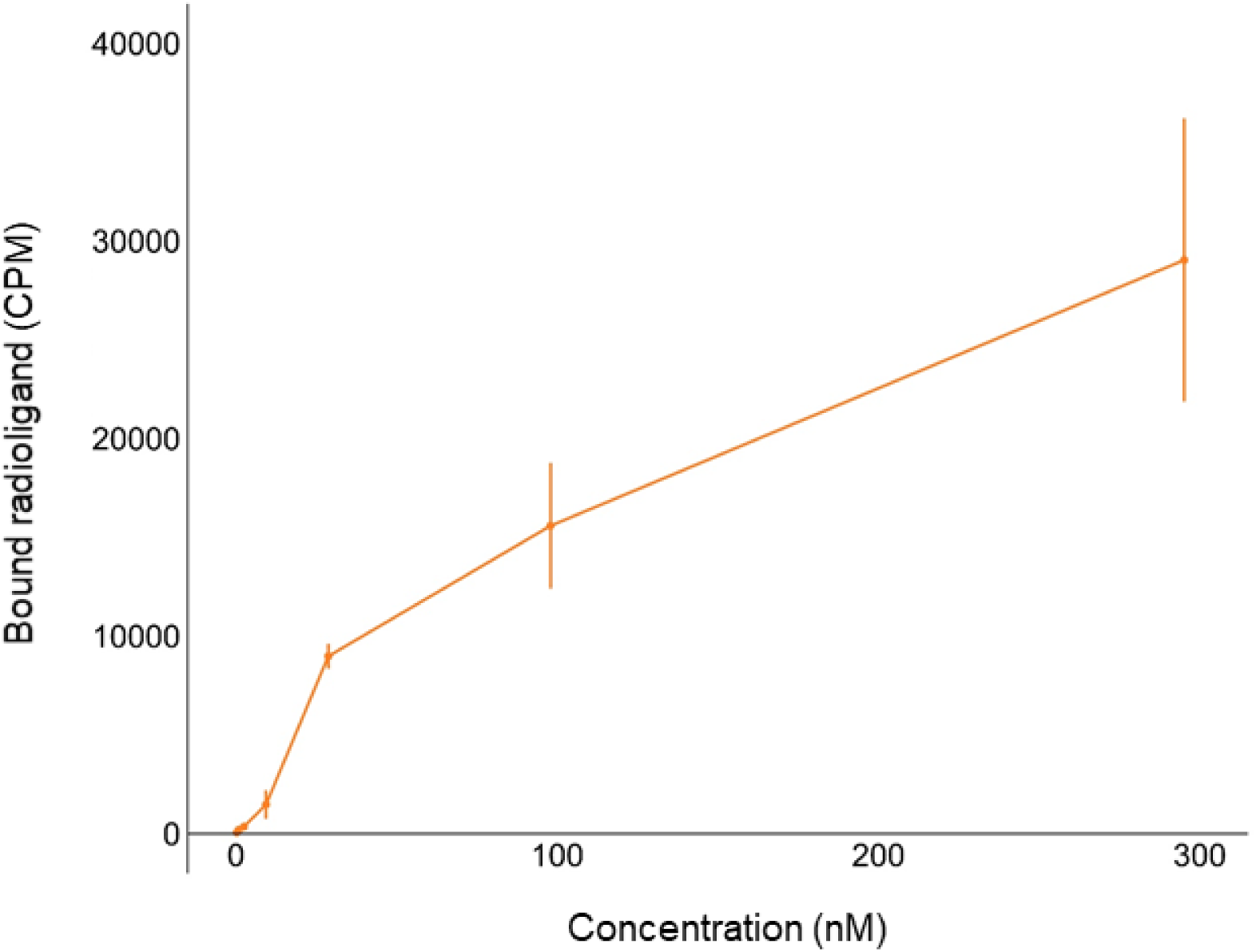
Saturation binding of agonist [^3^H]-dihydroalprenolol to membranes from HEK293 cells stably expressing β3AR-cc3.1. Saturation binding curve showing the specific binding of [^3^H]-dihydroalprenolol to β3AR-cc3.1. From the curve, a K_D_ value of 153 nM was estimated. n=3.

Although the ICL3 of most GPCRs is unstructured, our results are consistent with the observation that extended ICL3 structures are found in some natural GPCRs. For example, the ICL3 of squid rhodopsin [44] and bovine rhodopsin determined from its trigonal crystal[45] both form an extended anti-parallel helical structure that is similar to an antiparallel coiled coil [19]. These findings indicate that our engineered GPCR chimera might possibly even still bind G proteins and therefore be signaling-competent.

### β3AR-cc3.1 forms diffracting protein crystals

Because biophysical characterization of β3AR-cc3.1 demonstrated that our coiled-coil-based approach is well suited for quickly and efficiently stabilizing the GPCR, we next aimed at crystallizing the engineered thermostabilized variant bound to carvedilol. Initially, vapor diffusion at 22°C was employed using a protein concentration of 7.5mg/ml. Extensive crystallization trials resulted in the formation of 30-70μm-long needle-like crystals (Fig. 5A) of β3AR-cc3.1 in 0.2M MgCl_2_, 0.1M sodium cacodylate pH 6.5 and 50% v/v PEG 200. Typically, the crystals grew within two hours and reached their maximal size after 3-4 days. Crystals displayed diffraction to approximately 22Å (Fig. 5B). Intensive optimization was performed subsequently using different protein concentrations, additives, pHs, salts, buffers and precipitants, but there was no improvement in diffraction quality. We also tried to crystallize β3AR-cc3.1 using the lipidic cubic phase method [46], but did not obtain any crystals.

**Figure 5.**
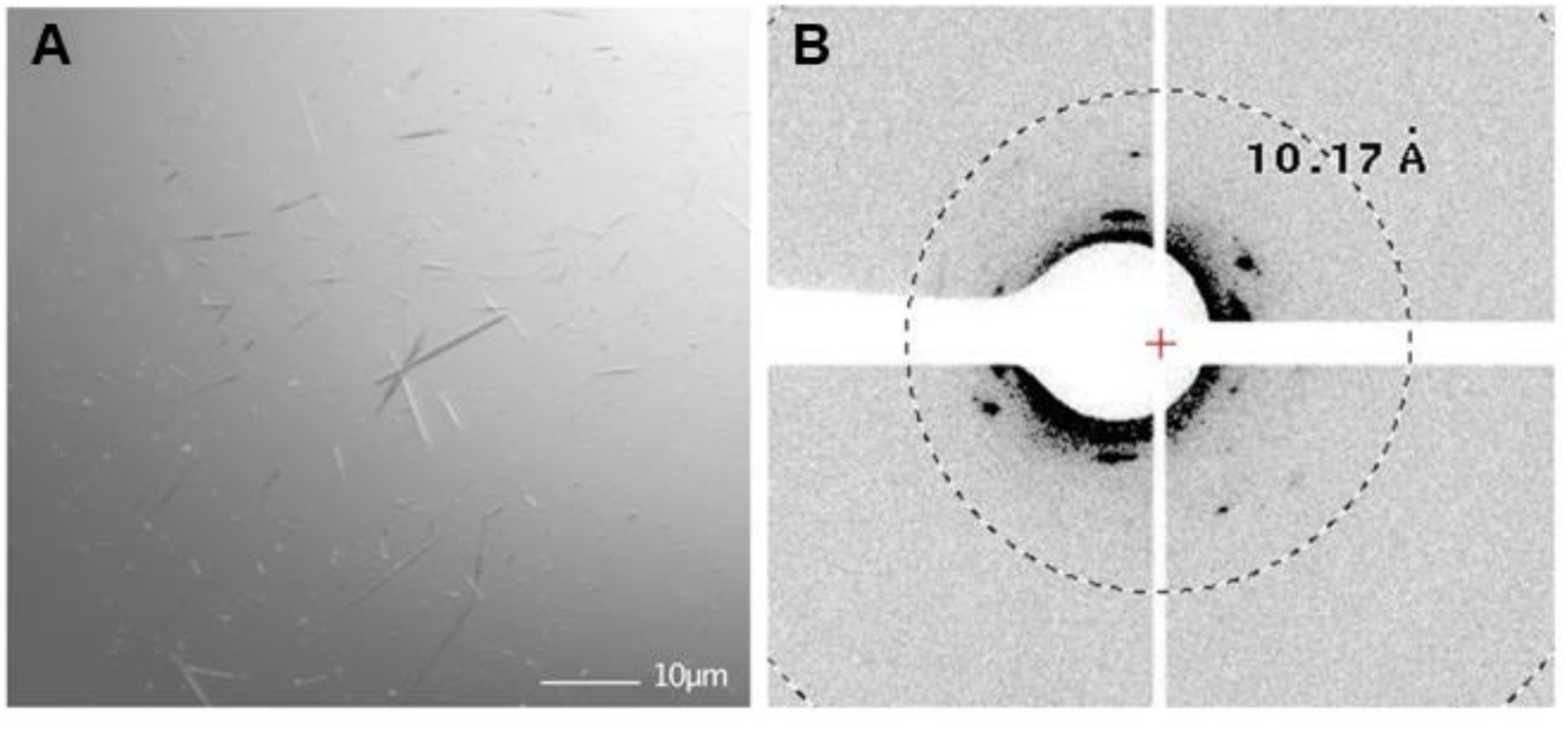
Crystallization of β3AR-cc3.1. (**A**) β3AR-cc3.1crystals. The needles grew at a protein concentration of 7.5mg in 0.2M MgCl_2_, 0.1M sodium cacodylate pH 6.5 pH and 50% v/v PEG 200 in 3-4 days. Size bar, 10µM (**B**) X-ray diffraction pattern of β3AR-cc3.1 crystals. The crystals diffracted to a resolution of about 22Å.

### Increasing crystal contacts by replacing the loop connecting the α-helices of the SRS coiled coil by T4L

Because we were not successful in optimizing conditions to obtain crystals that were suitable for structure determination, we decided to modify β3AR-cc3.1 on the protein level to improve crystal contacts. Truncation or mutation of the antiparallel coiled coil was not considered a valid option because the β3AR-cc3.1-SS and β3AR-cc3.2-SS variants failed to express, which was probably due to destabilization of the proteins. Although it has been reported that the loop connecting the two helices of an antiparallel coiled coil is crucial for the stability of the structure, it was demonstrated that substitution of the loop by a disulfide bond flanking the heptad repeats restored coiled-coil formation [30]. Although substitution of the connecting loop might alter the thermal stability of the antiparallel coiled coil, we decided to replace it with phage T4L. We selected T4L because it has been very successfully used as a fusion protein to crystallize and determine the structures of several GPCRs [9]. Because it crystallizes easily under many different conditions, T4L is considered an ideal fusion partner to establish crystal contacts. Another criterium for of choice was the existence of a β2AR crystal structure in which ICL3 was replaced by T4L. For the design, we used the structure-guided strategy described for the construction of β3AR-cc3.1. More specifically, we grafted the experimentally determined boundary of T4L to the coiled-coil like structure seen in β2AR on the antiparallel coiled coil of β3AR-cc3.1 (Fig. 6A and B).

**Figure 6.**
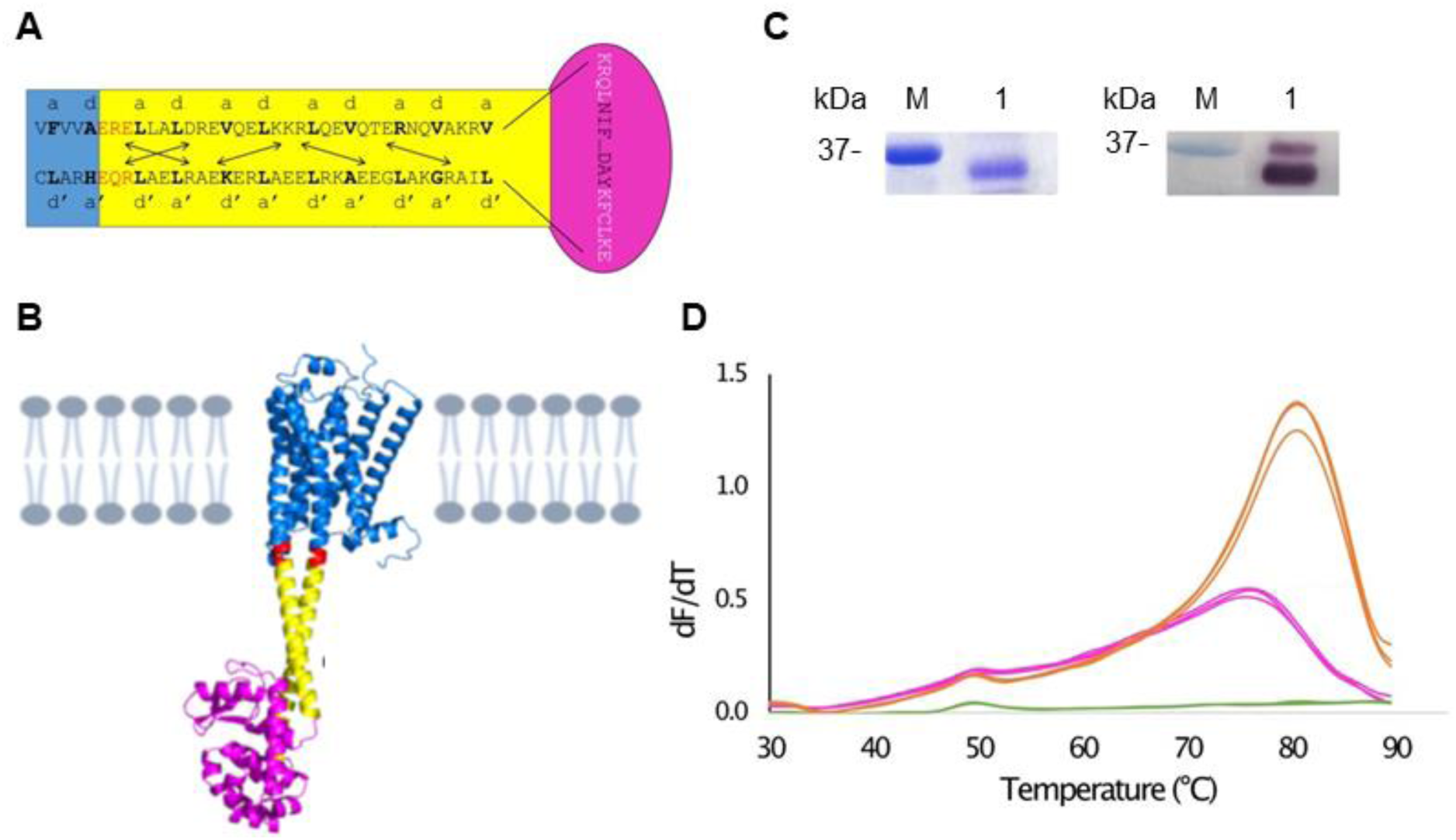
Construct design, purification and stability of β3AR-cc-T4L. **(A)** Schematic representation of β3AR-cc-T4L. For detail concerning the β3AR-cc3.1 part, see Fig. 1B. The T4L part is shown in magenta. The T4L sequence is in black and the connecting residues originating from β2AR are in white. (**B**) β3AR-cc-T4L model. β3AR is shown in blue, the cc3.1 coiled coil in yellow, T4L in magenta and residues at the junction in red. The model was generated by AlphaFold2 [7]. (**C**) SDS-PAGE (left panel) and western blot (right panel) analysis of purified β3AR-cc-T4L (lane 1). The migration of a marker protein (M) is shown. (**D**) Replicate CPM measurements of β3AR-cc3.1 (5 µg) and β3AR-cc-T4L (5 µg) bound to carvedilol in DDM. The T_m_ values of β3AR-cc3.1 (magenta) and β3AR-cc-T4L (orange) are 64.5 °C and 75 °C, respectively. The green curves represent measurements of the buffer without any protein.

Expression of the resulting variant, termed β3AR-cc-T4L, was only observed in T-REx-CHO cells (Fig. 6C). Although the protein showed two bands after Western blot analysis, we established stable T-REx-CHO cells expressing β3AR-cc-T4L.

Unfortunately, we were not able to adapt the cells for growth in suspension, we managed to purify a small amount of β3AR-cc-T4L for characterization from adherent T-REx-CHO cells using the best condition described for β3AR-cc3.1 (detergent combination DDM/CHS and antagonist carvediol). Notably, β3AR-cc-T4L was significantly more stable than β3AR-cc3.1 and yielded an almost 10.5°C higher T_m_ value of 75°C (Fig. 6D). Therefore, β3AR-cc-T4L represents a promising candidate for crystallization.

## Conclusions

Based on the available GPCR crystal structures, a combination of protein stabilization and accessible soluble surface to establish crystal contacts would represent an ideal tool for crystallization. Towards this aim, we identified thermostable antiparallel coiled coils a such a tool. Preplacement of ICL3 of β3AR by cc3.1 resulted in significant stabilization of the GPCR while retaining its ligand-binding properties. Furthermore, we were also able to stabilize two other GPCRs, 5HTR2C and α1BAR (not shown), demonstrating that this approach is generally suitable for the stabilization of GPCRs. Stabilized GPCRs should, for example, be of considerable interest for drug discovery applications where the wild-type protein is of limited stability.

Although we managed to crystallize the β3AR/coiled-coil chimera, the quality of the crystals even after extensive optimization was not suitable for structure determination. To supply additional surface for promoting crystal contacts, we replaced in a structure-based approach the loop connecting the helices of the antiparallel coiled coil by T4L. Although expression levels are currently not suitable for crystallization, we were able to show that the engineered GPCR is even more stable than the β3AR/coiled-coil chimera. To increase protein yields, we will screen β3AR proteins of different species. Notably, β3AR-cc-T4L could also become of interest for structure determination by cryo-EM. In a recent study using phase plates it was possible to reach 3.2Å resolution for streptavidin (52kDa) [47] that has a similar size as β3AR-cc-T4L.

## MATERIALS AND METHODS

### Constructs

Synthetic genes encoding the human β3 adrenergic receptor (β3AR) lacking amino-acid residues Pro3 to Thr27 and Pro369 to Ser408 and with the antiparallel coiled-coil sequences described in this study (table S1) inserted between Ala231 and Glu286 were codon-optimized for expression in human cells (Genewiz). The β3AR sequences also include the potentially thermostabilizing mutations Glu36Ala, Met86Val, Ile125Val, Glu126Trp, Tyr234Ala, Phe341Met, and Tyr346Leu [35]. Insert sequences were further modified to contain a hemaggutinin signal sequence followed by a modified FLAG tag at the N-terminus and a TwinStrep tag that can be removed by thrombin or HRV 3C cleavage at the C-terminus (Fig. S2). The full-length wild-type human β3AR cDNA sequence was used as a control. Insert sequences were subcloned into mammalian expression vectors pACMV-tetO [48] and pcDNA4/TO (ThermoFisher Scientific).

### Cell culture and protein expression

Human embryonic kidney HEK293T were used for transient small-scale expression tests. HEK293T, T-REx-293, T-REx-CHO (Thermo Fisher) and HEK293S GnTI^-^were used for stable expression. Cells were transiently or stably transfected using 25 kDa branched PEI as described [49]. The cells were grown adherently and maintained at 37°C in 5% CO_2_ in Dulbecco’s Modified Eagle Medium (DMEM) containing 10% fetal calf serum (FCS), penicillin (100U/ml), streptomycin (100μg/ml) and L-glutamine (2mM).

Stable cell lines were generated by using geneticin (G-418, 0.2mg/ml) or zeocin (0.4mg/mL) for pACMV-tetO and pcDNA4/TO, respectively. Individual colonies (24 for each receptor construct) typically appeared after 14 days and were isolated and expanded as described before [50]. Protein expression was induced with tetracycline (2μg/ml) and sodium butyrate (5 mM) and cells further incubated for 72 h. For each clone, the expression level of the recombinant protein was assessed by western blotting, and the best clones were selected for further large-scale expression in suspension. Stably transfected cells were grown in suspension in a final volume of 5×1l of PEM media (5% fetal calf serum (FCS), 1% (pyrrolidone carboxylic acid) PSA, 2.5mM Glutamax). Expression was induced upon reaching a cell density of ∼3×10^6^ cells/ml as described above. Cells were harvested after 72h by centrifugation at 2’500g, frozen in liquid nitrogen and stored at -80°C for protein purification or washed twice with PBS prior to freezing for membrane preparations.

### Membrane preparation and protein purification

For membrane preparation, cells were thawed on ice for 30min. The cells were lysed in 20mM Tris, pH 8, 500mM NaCl, 15% glycerol, 50mg/L DNAse I, 1 tablet/5L cell suspension of the EDTA-Free cOmplete Protease Inhibitor Cocktail (Roche) and 20μM carvedilol using a continuous flow EmulsiFlex-C3 cell disruptor (Avestin). The cell lysate was centrifuged at 10’000g for 30min at 4°C, followed by centrifugation of the resulting supernatant at 100’000g for 1h at 4°C. The membranes were flash-frozen in liquid nitrogen and stored at -80°C for further use.

For protein purification, membranes were thawed on ice and solubilized using 1% (w/v) n-dodecyl-β-d-maltopyranoside (DDM, Anatrace), 0.2% (w/v) cholesteryl hemisuccinate (CHS, Sigma-Aldrich) in 50mM Tris pH 7.5, 150mM NaCl containing 20μM carvedilol (TOCRIS) for 45min at 4°C on roller shaker. The insoluble material was separated by high-speed centrifugation at 100’000g for 1h at 4°C. The supernatant was loaded on a StrepTrap Sepharose High-Performance column (MERCK) equilibrated with 50mM Tris pH 7.8 and 150mM NaCl supplemented with 0.05% DDM, 0.01% CHS and 20μM carvedilol. The proteins were washed with 10 column volumes of 50mM Tris pH 7.8, 150mM NaCl, 0.05% DDM, 0.01% CHS, 20μM carvedilol and eluted with 4 column volumes of 50mM Tris pH 7.8, 150mM NaCl, 0.05% DDM, 0.01 % CHS, 20μM carvedilol and 2.5mM desthiobiotin. For size-exclusion chromatography (SEC), a HiLoad 10/30 Superdex-200 column (Life Sciences) was used. The buffer used for SEC was 50mM Tris pH 7.8, 150mM NaCl, 0.05% DDM and 0.02% CHS and 20μM carvedilol. The protein samples were concentrated to 30mg/ml using 100-kDa MWCO AmiconUltra concentrators (Millipore) for crystallization trials and further analysis.

### β3ARcc3.1 crystallization

β3ARcc3.1 bound to carvedilol was concentrated to 10 mg/ml and crystallized by sitting-drop vapor diffusion at 20°C using a TPP Mosquito robot. Proteins were mixed with the reservoir solution using a volume ratio of 1:1 (200nl each). Crystals of β3AR-cc3.1 were obtained in 0.2M MgCl_2_, 0.1M sodium cacodylate pH 6.5 and 50% v/v PEG 200. Crystals typically appeared within 4 hours and grew to their maximum size of 14 × 5 × 3μm within 2 days. Diffraction experiments performed at beamline PXI (Swiss Light Source, Villigen, Switzerland) equipped with an EIGER 16M high resolution diffractometer (Dectris) confirmed the existence of protein crystals. Subsequently, manual optimization of β3ARcc3.1 crystals was tried at a protein concentration of 15mg/ml.

### Radioligand binding assay

Membrane preparations ranged between 0.25-2µg of protein/well. The radioligand binding experiments were done in a volume of 200µl (50µl Hanks’ Balanced Salt Solution (HBSS) assay buffer, 20mM HEPES, 0.1% BSA, pH 7.4; 25µl antagonist or assay buffer (depending on assay type), 50µl membrane solution (final protein concentration 5µg), 50µl scintillation proximity assay (SPA) solution (Perkin Elmer), and 25µl of dihydroalprenolol hydrochloride, levo-[ring, propyl-^3^H(N)] (PerkinElmer). For saturation binding experiments, we used up to 300nM of [3H]-dihydroalprenolol. Different dilutions of the radioligand were prepared in assay buffer corresponding to a concentration range of approximately 0.02–300nM. Non-specific binding was determined in the presence of 10µM of the selective β3 antagonist L748337 (TOCRIS). Samples were incubated in 96-well plate sealed with transparent Topseal for 2h at 25°C with gentle agitation. Samples were centrifuged for 10min at 2’500g before being analyzed in a β-counter. Data were fitted as one-site binding using Prism. Statistical analyses were performed with Prism (GraphPad) using the unpaired t test.

### Thermostability assay

Thermal stability of proteins was assessed the fluorescent cysteine-reactive dye, 7-diethylamino-3-(4-maleimidophenyl)-4-methylcoumarin (CPM) as described before [40]. The protein concentration used per assay was 5-10 µg. Thermal unfolding was monitoring using the Qiagen Rotor-Gene Q instrument. Excitation was at 365 nm and emission at 460 nm was recorded over a temperature range from 25 to 90 °C with a ramping rate of 2 °C/min. Data analysis was performed using the Rotor-Gene software.

## ACKNOWLEDGMENTS

The PXI beamline staff (Swiss Light Source, Villigen, Switzerland) is acknowledged for their support during the diffraction experiments. We thank Mara Wieser and Shahrooz Nasrollahi Shirazi for technical assistance.

## AUTHOR CONTRIBUTIONS

R.A.K. conceptualization; M.A. and R.A.K. data curation; M.A., M.W., D.F. and X.L. formal analysis, investigation and validation; R.A.K. and X.L. funding acquisition and supervision; M.A. and R.A.K. writing-review and editing and writing original draft with input from the other authors.

## FUNDING AND ADDITIONAL INFORMATION

This work was supported by grants 31003A_163449 and 31003A_170028 of the Swiss National Science Foundation to R.A.K. and X.L., respectively.

