## Supplementary Information for "A Coiled-Coil-Based Design Strategy for the Thermostabilization of G-Protein-Coupled Receptors"

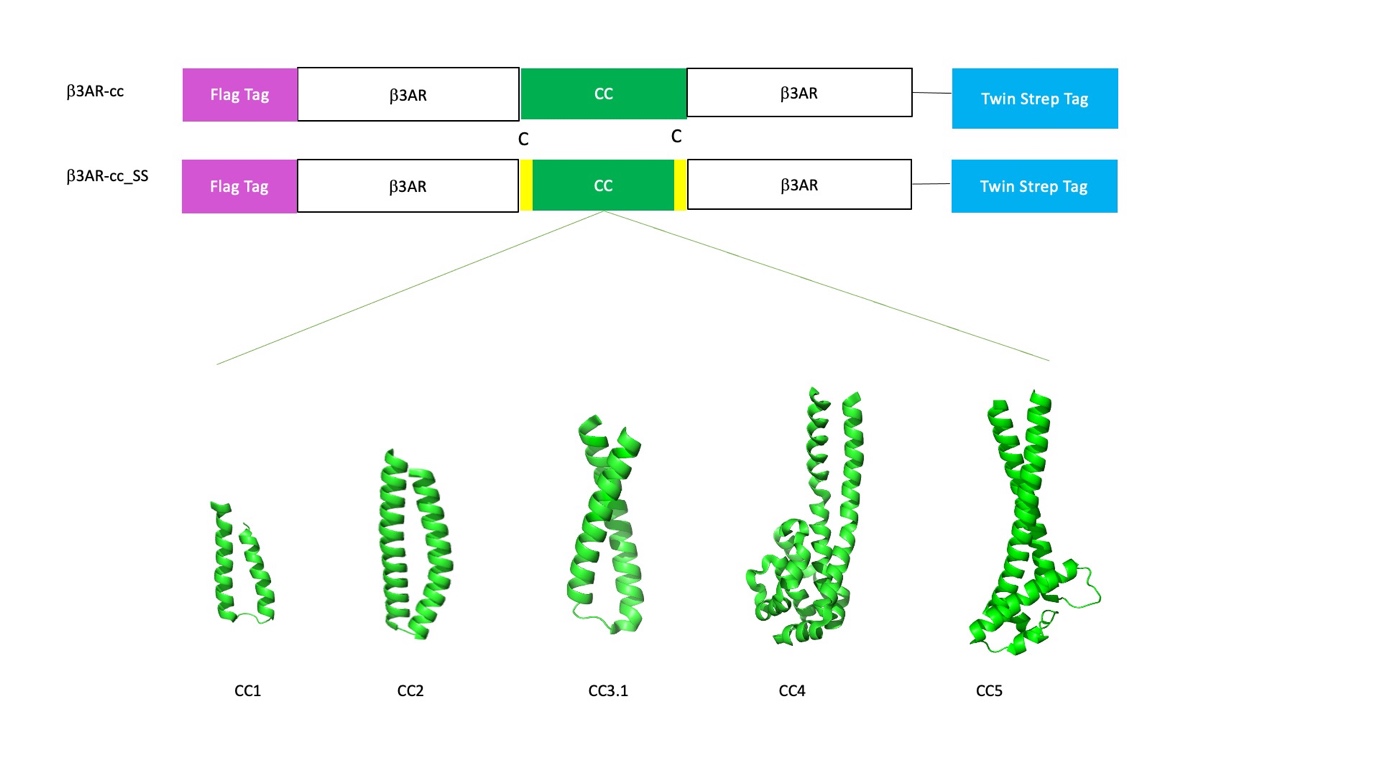

**Supplementary figure 1.** **Design of β3AR-cc constructs.** Schematic representation of the chimeric β3AR variants and crystal structures of coiled-coil domains that were used for replacing ICL3 of β3AR. For cc3.2 there is no structure available. The constructs have N-terminal Flag tag and C-terminal TwinStrep tag for western blot detection and protein purification.

**Supplementary table 1.**

| Coiled-coil | Sequence | PDB ID | Resolution |
| --- | --- | --- | --- |
| cc1 | CKIEHAKKKRLFDLYINGSYEVSELDSMMNDIDAQINYC | 5UDO | 2.5 Å |
| cc2 | LKEVQDNITLHEQRLVTTRQKLKDAERAVELDPDDVNKSTLQSRRAAVSALETKLGELKRELADL | 2IC6 | 1.15 Å |
| cc3.1 | LLALDREVQELKKRLQEVQTERNQVAKRVPKAPPEEKEALIARGKALGEEAKRLEEALREKEARLEAL | 1SRY | 2.5 Å |
| cc3.2 | VRELDELWRKKLQEVNEVRHKHNVVTRMIAKARDPEERKRLIEEARRLLKLREELEKELKRIEEEREKL | NA | NA |
| cc4 | LKEEQERKAEIQADIAQQEKNKAKLVVDRNKIIESQDVIRQYNLADMFKDYIPNISDLDKLDLANPKKELIKQAIKQGVEIAKKILGNISKGLKYIELADARAKLDERINQINKDCDDLKIQLKGVEQRIAGI | 6EK4 | 2.8 Å |
| cc5 | LLALLAVDEQLHKQQEVIADKQMSVKEDLDKVEPAVIEAQNAVKSIKKQHLVEVRSMANPPAAVKLALESIALLLGESTTDWKQIRSIIMRENFIPTIVNFSAEEISDAIREKMKKNYMSNPSYNYEIVNRASLAAGPMVKWAIAQLNYADMLKRVEPLRNELQKLEDDAKDNQQKLEAL | 3ERR | 2.2 Å |
| cc-T4L | LLALDREVQELKKRLQEVQTERNQVAKRVKRQLNIFEMLRIDEGLRLKIYKDTEGYYTIGIGHLLTKSPSLNAAKSELDKAIGRNTNGVITKDEAEKLFNQDVDAAVRGILRNAKLKPVYDSLDAVRRAALINMVFQMGETGVAGFTNSLRMLQQKRWDEAAVNLAKSRWYNQTPNRAKRVITTFRTGTWDAYKFCLKELIARGKALGEEAKRLEEALREKEARLEAL | NA | NA |

Amino acid sequences of the coiled-coil sequences used in this study. The underlined amino acids were changed to Cys in the disulfide bridge variants.
